# The salivary and nasopharyngeal microbiomes are associated with SARS-CoV-2 infection and disease severity

**DOI:** 10.1101/2022.05.31.494162

**Authors:** Josh G. Kim, Ai Zhang, Adriana M. Rauseo, Charles W. Goss, Philip A. Mudd, Jane A. O’Halloran, Leyao Wang

## Abstract

Oral and upper respiratory microbiota play important roles in modulating host immune responses to viral infection. As emerging evidence suggests the host microbiome may be involved in the pathophysiology of COVID-19, we aimed to investigate associations between the oral and nasopharyngeal microbiome and COVID-19 severity. We collected saliva (*n* = 78) and nasopharyngeal swab (*n* = 66) samples from a COVID-19 cohort and characterized the microbiomes using 16S ribosomal RNA gene sequencing. We also examined associations between the salivary and nasopharyngeal microbiome and age, COVID-19 symptoms, and blood cytokines. SARS-CoV-2 infection status, but not COVID-19 severity, was associated with community-level differences in the oral and nasopharyngeal microbiomes. Salivary and nasopharyngeal microbiome alpha diversity negatively correlated with age and were associated with fever and diarrhea. Several bacterial genera were differentially abundant by COVID-19 severity, including oral *Bifidobacterium, Lactobacillus*, and *Solobacterium*, all of which were depleted in patients with severe COVID-19. Nasopharyngeal *Paracoccus* was depleted while nasopharyngeal *Proteus, Cupravidus, and Lactobacillus* were increased in patients with severe COVID-19. Further analysis revealed that the abundance of oral *Bifidobacterium* was negatively associated with plasma concentrations of known COVID-19 biomarkers interleukin 17F (IL-17F) and monocyte chemoattractant protein-1 (MCP-1). In conclusion, our results suggest COVID-19 disease severity is associated with the relative abundance of certain bacterial taxa.

## Introduction

Coronavirus disease 2019 (COVID-19), caused by severe acute respiratory syndrome coronavirus 2 (SARS-CoV-2), is a global public health crisis. As of May 2022, SARS-CoV-2 has infected over 528,000,000 people and caused over 6,200,000 deaths worldwide according to the Johns Hopkins coronavirus resource center. A particularly challenging feature of the COVID-19 pandemic has been the extremely wide range of disease severity experienced by infected individuals. While SARS-CoV-2 infection may cause only asymptomatic carriage or mild symptoms in some individuals, it can result in severe lung damage or death in others. Therefore, the identification of early biomarkers that can infer COVID-19 disease severity is critical.

Accumulating evidence suggests that the oral cavity is a robust portal for SARS-CoV-2 entry, replication, and shedding. Host factors important for SARS-CoV-2 entry, including angiotensin converting enzyme 2 (ACE2) and serine protease TMPRSS family members (TMPRSS2 and TMPRSS4), are highly expressed in oral epithelial cells and salivary glands (1-4). Viral infection in the oropharynx is likely to be influenced by the human oral microbiota, which is the second largest and most diverse microbiota in the human body, with over 700 species of bacteria that play major roles in maintaining local homeostasis and modulating immune responses towards invading pathogens (5-7). A growing body of evidence points toward the role of the oral microbiome in the establishment and progression of SARS-CoV-2 infection. For instance, patients with COVID-19 have significantly disrupted oropharyngeal microbiomes compared to patients with flu or healthy individuals (8-10). Oral microbial dysbiosis was associated with severe symptoms of COVID-19, increased local inflammation, duration of COVID-19 symptoms, and more recently, long COVID (11-13). In addition, some elevated bacterial taxa correlated with systemic inflammatory markers such as a high neutrophil-lymphocyte ratio, suggesting that the oral microbiota may be a sensitive biomarker or even play a role in the activation or suppression of innate and humoral immunity against SARS-CoV-2 infection (8).

The nasopharyngeal microbiota is also of interest because the nasal mucosa is a major site of SARS-CoV-2 viral infection, replication, and dissemination in the host (14). Infection of the nasal mucosal surfaces occurs in the context of the nasopharyngeal microbiota, which plays a major role in mucosal homeostasis and progression of viral infections (15). On one hand, viral infection may lead to bacterial co-infection, a major cause of mortality in previous viral pandemics such as the 2009 H1N1 influenza outbreak (16). On the other hand, pre-existing microbial dysbiosis could induce skewed inflammatory responses during respiratory viral infections and lead to increased risk of severe outcomes (17). A study analyzing oropharyngeal, nasopharyngeal, and endotracheal samples from hospitalized COVID-19 patients using 16S rRNA sequencing reported respiratory tract bacterial dysbiosis in patients with COVID-19; microbial signatures were also associated with COVID-19 severity and systemic immune response (12). Recent studies have also reported lower airway microbial signatures associated with poor clinical outcome of COVID-19 and upper respiratory microbiota associated with mortality and COVID-19 severity (18, 19). These early findings suggest that the airway microbiome may be an important factor in indicating and influencing COVID-19 clinical outcomes and should be investigated further.

Given the importance of the host microbiome in indicating and mediating immune responses to respiratory viral infections, we hypothesized that the salivary and nasopharyngeal microbiomes are associated with COVID-19 disease severity. To test this hypothesis, we collected saliva and nasopharyngeal swab samples from a well-characterized COVID-19 cohort and extracted microbial DNA for 16S ribosomal RNA (rRNA) gene sequencing to identify salivary and nasopharyngeal microbial features associated with COVID-19 severity.

## Results

### 1. Patients with SARS-CoV-2 infection harbored significant compositional differences in the salivary and nasopharyngeal microbiome compared to patients without SARS-CoV-2

We collected saliva (*n* = 78) and nasopharyngeal swab (*n* = 66) samples from patients who presented for SARS-CoV-2 testing with symptoms consistent with COVID-19. The saliva samples included 60 from SARS-CoV-2-positive patients and 18 from SARS-CoV-2-negative patients, and the nasopharyngeal swab samples included 54 from SARS-CoV-2-positive patients and 12 from SARS-CoV-2-negative patients. Demographics and clinical outcomes of this study population, stratified by COVID-19 status, are shown in **Table 1** for saliva samples and **Table 2** for nasopharyngeal swab samples. For patients from whom saliva samples were collected, age was significantly higher in individuals who tested positive for SARS-CoV-2 (*P* = 0.03). Congestive heart failure was also significantly higher in the SARS-CoV-2-positive group (*P* = 0.03). For patients from whom nasopharyngeal swab samples were collected, significantly more patients who tested positive for SARS-CoV-2 were of African American race (*P* = 0.001). For both saliva and nasopharyngeal swab samples, significantly more SARS-CoV-2-positive patients had diabetes and were hospitalized than SARS-CoV-2-negative patients. There were no other significant differences in other baseline characteristics such as sex, BMI, current smoking status, chronic pulmonary disease, and obesity.

**Table 1.** Patient characteristics for saliva samples

| Variable | SARS-CoV-2<br>negative (n = 18) | SARS-CoV-2<br>positive (n = 60) | P value |
| --- | --- | --- | --- |
| Age (years), mean (SD) | 44.85 (16.31) | 54.51 (16.40) | 0.03 |
| Sex: Male, n (%) | 10 (55.56) | 40 (66.67) | 0.561 |
| BMI, mean (SD) | 30.4 (9.73)<sup>b</sup> | 34.43 (10.87) | 0.146 |
| Race: African American, n (%) | 9 (50) | 40 (66.67) | 0.315 |
| Current smoker: Yes, n (%) | 4 (22.22) | 6 (10) | 0.227<sup>a</sup> |
| Hospitalization, n (%) | 2 (11.11) | 57 (95) | <.001<sup>a</sup> |
| Ventilator, n (%) | 0 (0) | 7 (11.67) | 0.192<sup>a</sup> |
| Death due to COVID, n (%) | 0 (0) | 3 (5) | 1<sup>a</sup> |
| Chronic pulmonary disease: Yes, n (%)<sup>c</sup> | 5 (35.71) | 16 (30.77) | 0.724 |
| Congestive heart failure: Yes, n (%)<sup>c</sup> | 0 (0) | 14 (26.92) | 0.03<sup>a</sup> |
| Diabetes: Yes, n (%)<sup>c</sup> | 3 (21.43) | 30 (57.69) | 0.033<sup>a</sup> |
| Obesity: Yes, n (%)<sup>c</sup> | 1 (7.14) | 15 (28.85) | 0.159<sup>a</sup> |
| Hypertension: Yes, n (%)<sup>c</sup> | 5 (35.71) | 31 (59.62) | 0.111 |

Significance was evaluated on the basis of the Mann-Whitney U test and the chi-squared test or Fisher's exact test. <sup>a</sup>P value calculated using Fisher's exact test. <sup>b</sup>One missing value. <sup>c</sup>Four missing values in SARS-CoV-2-negative group and eight missing values in SARS-CoV-2-positive group

**Table 2.** Patient characteristics for nasopharyngeal swab samples

| Variable | SARS-CoV-2<br>negative (n = 12) | SARS-CoV-2<br>positive (n = 54) | P value |
| --- | --- | --- | --- |
| Age (years), mean (SD) | 57.63 (17.87) | 64.34 (13.68) | 0.241 |
| Sex: Male, n (%) | 4 (33.33) | 30 (55.56) | 0.283 |
| BMI, mean (SD) | 26.58 (7.49) | 29.57 (8.08) | 0.178 |
| Race: African American, n (%) | 3 (25) | 42 (77.78) | 0.001<sup>a</sup> |
| Current smoker: Yes, n (%) | 0 (0) | 7 (12.96) | 0.330<sup>a</sup> |
| Hospitalization, n (%) | 2 (16.67) | 54 (100) | <.001<sup>a</sup> |
| Ventilator, n (%) | 1 (8.33) | 16 (29.63) | 0.163<sup>a</sup> |
| Death due to COVID, n (%) | 0 (0) | 6 (11.11) | 0.582<sup>a</sup> |
| Chronic pulmonary disease: Yes, n (%)<sup>b</sup> | 1 (10) | 15 (31.91) | 0.253<sup>a</sup> |
| Congestive heart failure: Yes, n (%)<sup>b</sup> | 0 (0) | 12 (25.53) | 0.1<sup>a</sup> |
| Diabetes: Yes, n (%)<sup>b</sup> | 2 (20) | 29 (61.7) | 0.032<sup>a</sup> |
| Obesity: Yes, n (%)<sup>b</sup> | 2 (20) | 6 (12.77) | 0.619<sup>a</sup> |
| Hypertension: Yes, n (%)<sup>b</sup> | 4 (40) | 34 (72.34) | 0.069<sup>a</sup> |

Significance was evaluated on the basis of the Mann-Whitney U test and the chi-squared test or Fisher's exact test. <sup>a</sup>P value calculated using Fisher's exact test. <sup>b</sup>Two missing values in SARS-CoV-2-negative group and seven missing values in SARS-CoV-2-positive group.

We profiled the salivary and nasopharyngeal microbiome by 16S rRNA gene sequencing. After quality filtering, the total number of reads for saliva samples was 1,389,970, and the mean number of reads was 17,820 per subject. For nasopharyngeal swab samples, the total number of reads was 563,829 and the mean was 8,543 per subject. The top five abundant phyla in the salivary microbiome were Firmicutes, Actinobacteria, Bacteroidetes, Proteobacteria, and Fusobacteria. Compared to patients without COVID-19, patients with COVID-19 harbored a reduced abundance of Proteobacteria and Fusobacteria in the salivary microbiome (**Figure 1A**). The top five abundant phyla in the nasopharyngeal microbiome were Firmicutes, Actinobacteria, Proteobacteria, Bacteroidetes, and Fusobacteria, and there were no significant differences in their abundances between patients infected with SARS-CoV-2 and patients not infected with SARS-CoV-2. (**Figure 1B**).

**Figure 1.**
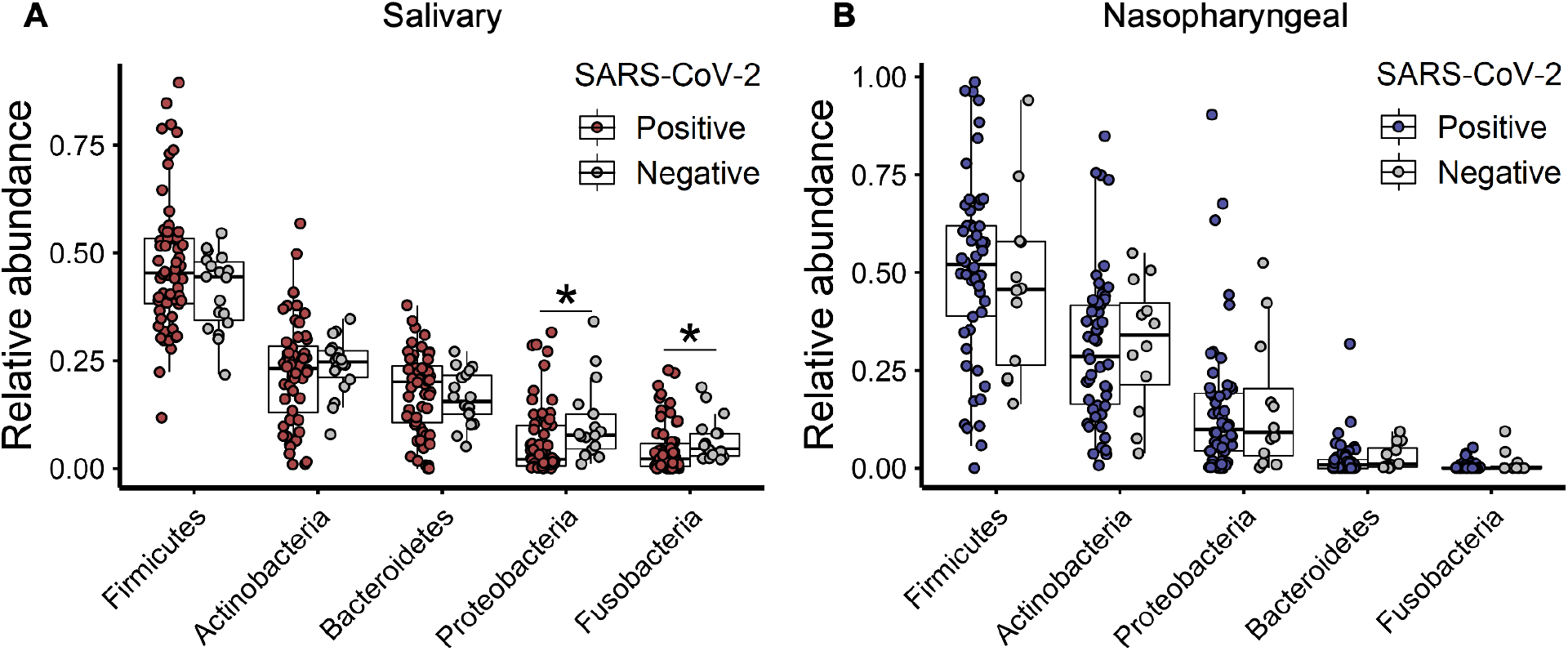
Relative abundances of the top five most abundant bacterial phyla in the salivary (A) and nasopharyngeal (B) microbial communities of SARS-CoV-2-positive subjects and SARS-CoV-2-negative controls. Statistical significance was assessed using Wilcoxon signed-rank test and analysis was adjusted for multiple comparisons using the Benjamini-Hochberg method. *, *P* < 0.05; **, *P* < 0.01.

To assess community level alterations of the salivary and nasopharyngeal microbiomes during SARS-CoV-2 infection, we compared alpha diversity and beta diversity between COVID-19 and non-COVID-19 patients. We observed a significant decrease in alpha diversity of salivary microbial communities of SARS-CoV-2-positive patients compared to SARS-CoV-2-negative patients (**Figure 2A**). Similarly, alpha diversity was significantly reduced in the nasopharyngeal microbiome of SARS-CoV-2-positive patients compared to SARS-CoV-2-negative patients, but only the richness index was significant (**Figure 2B**). Both salivary and nasopharyngeal microbial communities of SARS-CoV-2-positive patients differed markedly from those of SARS-CoV-2-negative patients based on principal coordinates analysis (PCoA) of UniFrac distances (**Figure 2C** and **Figure 2D**). In nasopharyngeal samples, this difference was only significant based on principal coordinates analysis of unweighted UniFrac distances (**Figure 2D**). The microbiome differences we observed between COVID-19-positive and COVID-19-negative subjects are consistent with previous studies on the oral and airway microbiome in COVID-19 (20-23). As there were some substantial differences in baseline characteristics between the two groups, we also compared the salivary and nasopharyngeal microbiome between selected sex-, age-, and race-matched SARS-CoV-2 positive (n = 18 for saliva samples, n = 9 for nasopharyngeal swab samples) and SARS-CoV-2 negative (n = 18 for saliva samples, n = 9 for nasopharyngeal swab samples) patients to validate our findings (**Figure S1**). We observed similar results for saliva samples, but no significant differences between SARS-CoV-2-positive and SARS-CoV-2-negative patients for nasopharyngeal swab samples, which may be due to the low number of samples available in the matched case-control group.

**Figure 2.**
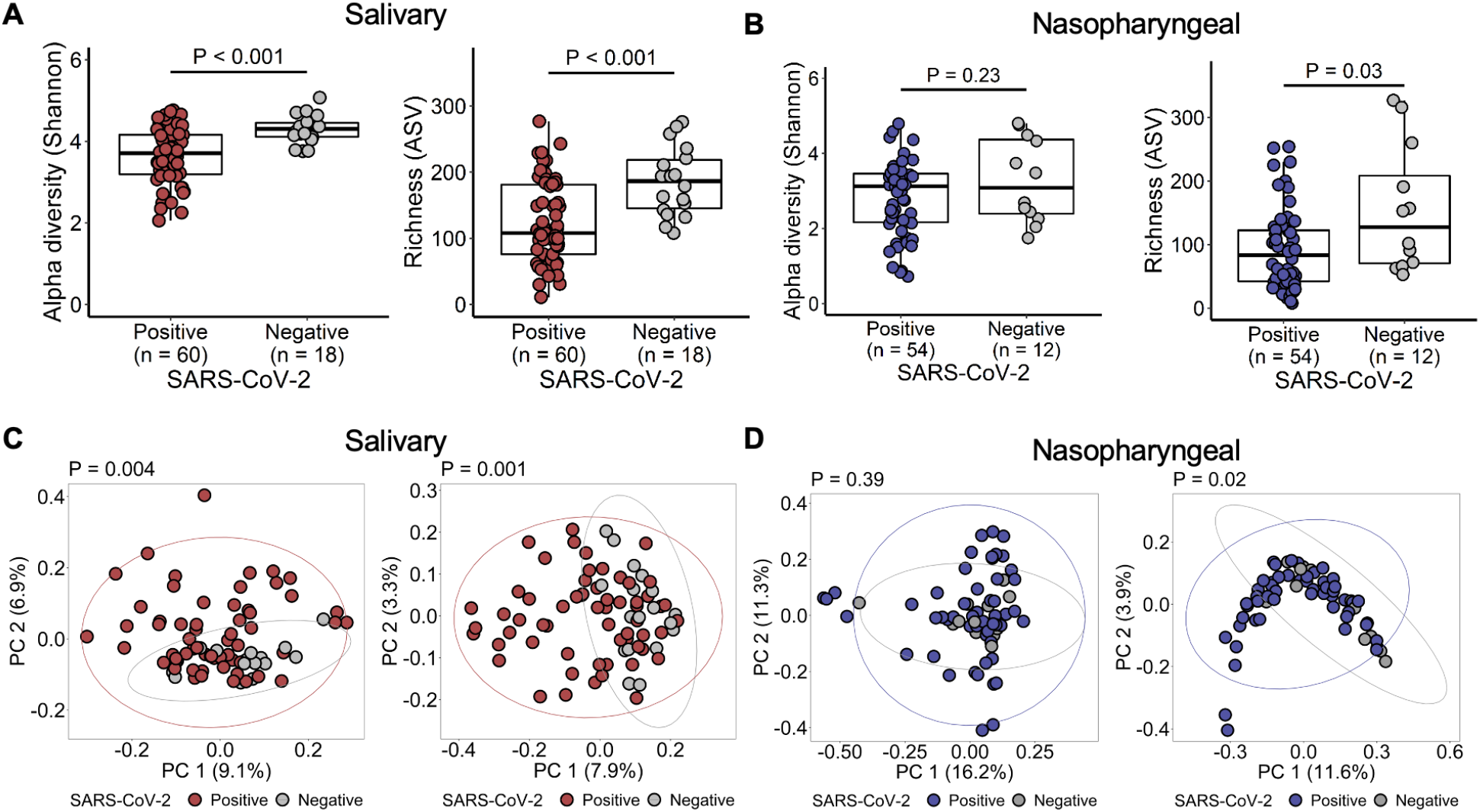
Salivary and nasopharyngeal microbial dysbiosis in COVID-19 patients. Alpha diversity of salivary (A) and nasopharyngeal (B) microbiomes of SARS-CoV-2-positive and SARS-CoV-2-negative patients. Principal coordinates analysis of weighted (C, D, left) and unweighted (C, D, right) UniFrac distances of salivary (C) and nasopharyngeal (D) microbial communities of SARS-CoV-2-positive and SARS-CoV-2-negative subjects. Statistical significance was assessed using Wilcoxon signed-rank test for panels A and B, and using PERMANOVA for panels C and D.

### 2. No compositional differences in the salivary and nasopharyngeal microbiome between COVID-19 patients who were later admitted to ICU and those who were not. Salivary and nasopharyngeal microbial communities are associated with COVID-19 symptoms and age

To investigate microbial features associated with severe outcomes of COVID-19, we then focused on SARS-CoV-2-positive patients (*n* = 60 for saliva samples, *n* = 54 for nasopharyngeal samples) and stratified them by COVID-19 severity according to intensive care unit (ICU) admission status. Demographics, symptoms, and clinical outcomes of the SARS-CoV-2-positive patients, stratified by ICU admission, are presented in **Table 3** and **Table 4**. For saliva samples, 18 patients were admitted to an ICU and 42 were not. Of the 18 ICU patients, 7 (38.89%) required mechanical ventilation and 3 (16.67%) died; no subjects in the non-ICU group died. The percentage of African American patients was higher in the non-ICU group than the ICU group (*P* = 0.036). For nasopharyngeal samples, 30 were admitted to an ICU and 24 were not. Of the 30 ICU patients, 16 (53.33%) required mechanical ventilation and 5 (16.67%) died; one (4.17%) subject in the non-ICU group died. We did not observe a difference in alpha diversity between ICU and non-ICU groups in the salivary microbiome nor the nasopharyngeal microbiome (**Figure S2A, Figure S2B**). The salivary and nasopharyngeal microbial compositions in the ICU group were not significantly different from those of the non-ICU group (**Figure S2C, Figure S2D**).

**Table 3.**
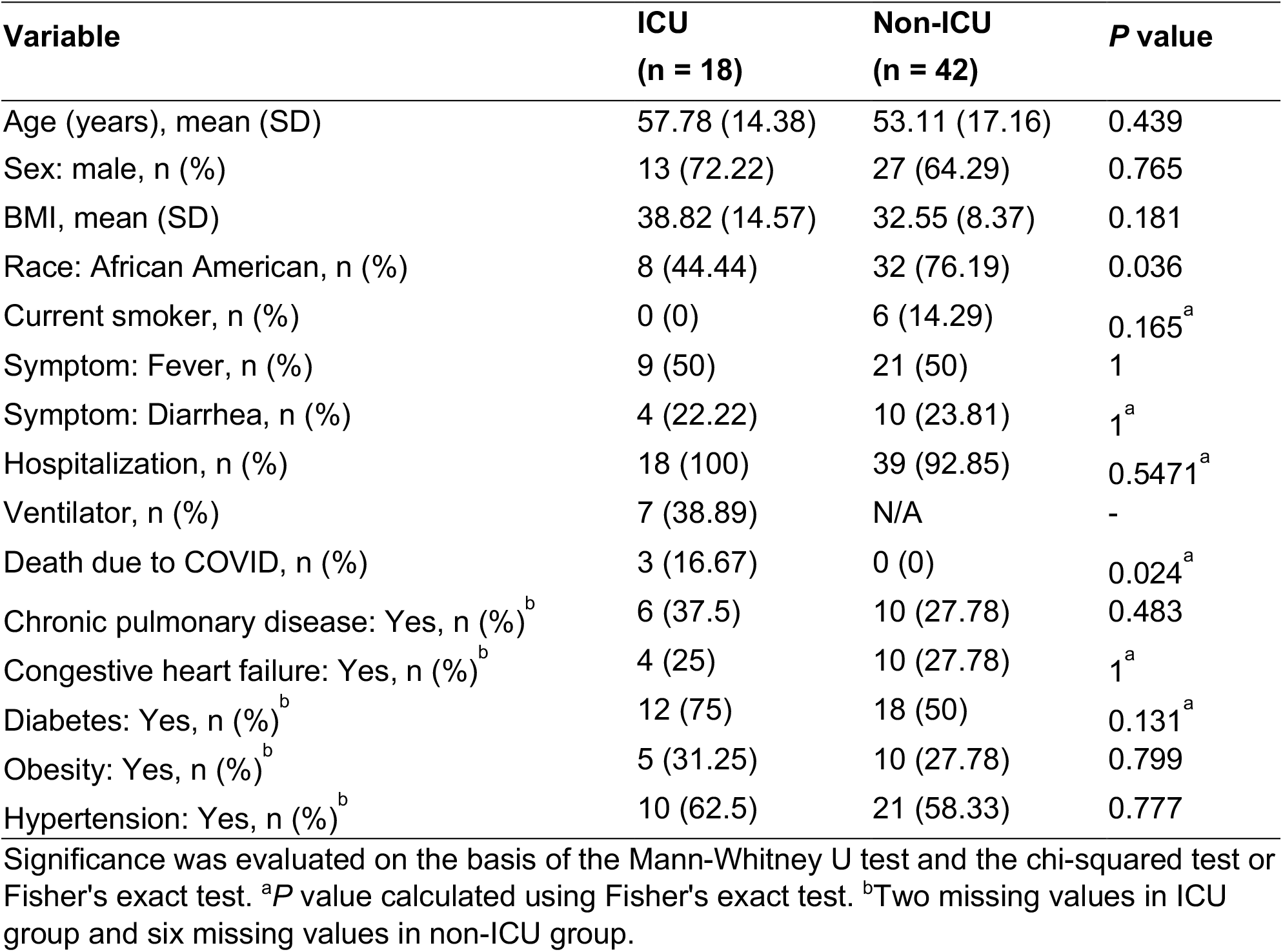
Patient characteristics of SARS-CoV-2-positive patients for saliva samples, stratified by ICU admission

| Variable | ICU<br>(n = 18) | Non-ICU<br>(n = 42) | P value |
| --- | --- | --- | --- |
| Age (years), mean (SD) | 57.78 (14.38) | 53.11 (17.16) | 0.439 |
| Sex: male, n (%) | 13 (72.22) | 27 (64.29) | 0.765 |
| BMI, mean (SD) | 38.82 (14.57) | 32.55 (8.37) | 0.181 |
| Race: African American, n (%) | 8 (44.44) | 32 (76.19) | 0.036 |
| Current smoker, n (%) | 0 (0) | 6 (14.29) | 0.165<sup>a</sup> |
| Symptom: Fever, n (%) | 9 (50) | 21 (50) | 1 |
| Symptom: Diarrhea, n (%) | 4 (22.22) | 10 (23.81) | 1<sup>a</sup> |
| Hospitalization, n (%) | 18 (100) | 39 (92.85) | 0.5471<sup>a</sup> |
| Ventilator, n (%) | 7 (38.89) | N/A | - |
| Death due to COVID, n (%) | 3 (16.67) | 0 (0) | 0.024<sup>a</sup> |
| Chronic pulmonary disease: Yes, n (%)<sup>b</sup> | 6 (37.5) | 10 (27.78) | 0.483 |
| Congestive heart failure: Yes, n (%)<sup>b</sup> | 4 (25) | 10 (27.78) | 1<sup>a</sup> |
| Diabetes: Yes, n (%)<sup>b</sup> | 12 (75) | 18 (50) | 0.131<sup>a</sup> |
| Obesity: Yes, n (%)<sup>b</sup> | 5 (31.25) | 10 (27.78) | 0.799 |
| Hypertension: Yes, n (%)<sup>b</sup> | 10 (62.5) | 21 (58.33) | 0.777 |

Significance was evaluated on the basis of the Mann-Whitney U test and the chi-squared test or Fisher's exact test. <sup>a</sup>P value calculated using Fisher's exact test. <sup>b</sup>Two missing values in ICU group and six missing values in non-ICU group.

**Table 4.** Patient characteristics of SARS-CoV-2-positive patients for nasopharyngeal swab samples, stratified by ICU admission

| Variable | ICU<br>(n = 30) | Non-ICU<br>(n = 24) | P value |
| --- | --- | --- | --- |
| Age (years), mean (SD) | 66.54 (11.81) | 61.60 (15.53) | 0.21 |
| Sex: male, n (%) | 17 (56.67) | 13 (54.17) | 0.927 |
| BMI, mean (SD) | 27.93 (7.34) | 31.61 (8.64) | 0.077 |
| Race: African American, n (%) | 23 (76.67) | 19 (79.17) | 0.913 |
| Current smoker, n (%) | 3 (10) | 4 (16.67) | 0.687<sup>a</sup> |
| Symptom: Fever, n (%) | 10 (33.33) | 11 (45.83) | 0.512 |
| Symptom: Diarrhea, n (%) | 1 (3.33) | 4 (16.67) | 0.159<sup>a</sup> |
| Hospitalization, n (%) | 30 (100) | 24 (100) | - |
| Ventilator, n (%) | 16 (53.33) | N/A | - |
| Death due to COVID, n (%) | 5 (16.67) | 1 (4.17) | 0.21<sup>a</sup> |
| Chronic pulmonary disease: Yes, n (%)<sup>b</sup> | 9 (34.62) | 6 (28.57) | 0.659 |
| Congestive heart failure: Yes, n (%)<sup>b</sup> | 4 (15.38) | 8 (38.1) | 0.1<sup>a</sup> |
| Diabetes: Yes, n (%)<sup>b</sup> | 18 (69.23) | 11 (52.38) | 0.237 |
| Obesity: Yes, n (%)<sup>b</sup> | 1 (3.85) | 5 (23.81) | 0.076<sup>a</sup> |
| Hypertension: Yes, n (%)<sup>b</sup> | 19 (73.08) | 15 (71.43) | 0.9 |

Significance was evaluated on the basis of the Mann-Whitney U test and the chi-squared test or Fisher's exact test. <sup>a</sup>P value calculated using Fisher's exact test. <sup>b</sup>Four missing values in ICU group and three missing values in non-ICU group.

We also investigated whether community-level microbial alterations were associated with several major symptoms of COVID-19 including fever, coughing, shortness of breath, diarrhea, and nausea/vomiting, by comparing alpha and beta diversity of the salivary or nasopharyngeal microbiome in patients with or without these symptoms. We observed significantly greater salivary microbiota alpha diversity in COVID-19 patients reporting diarrhea compared to those not reporting diarrhea (**Figure 3A**). In addition, alpha diversity of the nasopharyngeal microbiota was reduced in patients with fever compared to those without fever, though this was only significant for richness (**Figure 3A**).

**Figure 3.**
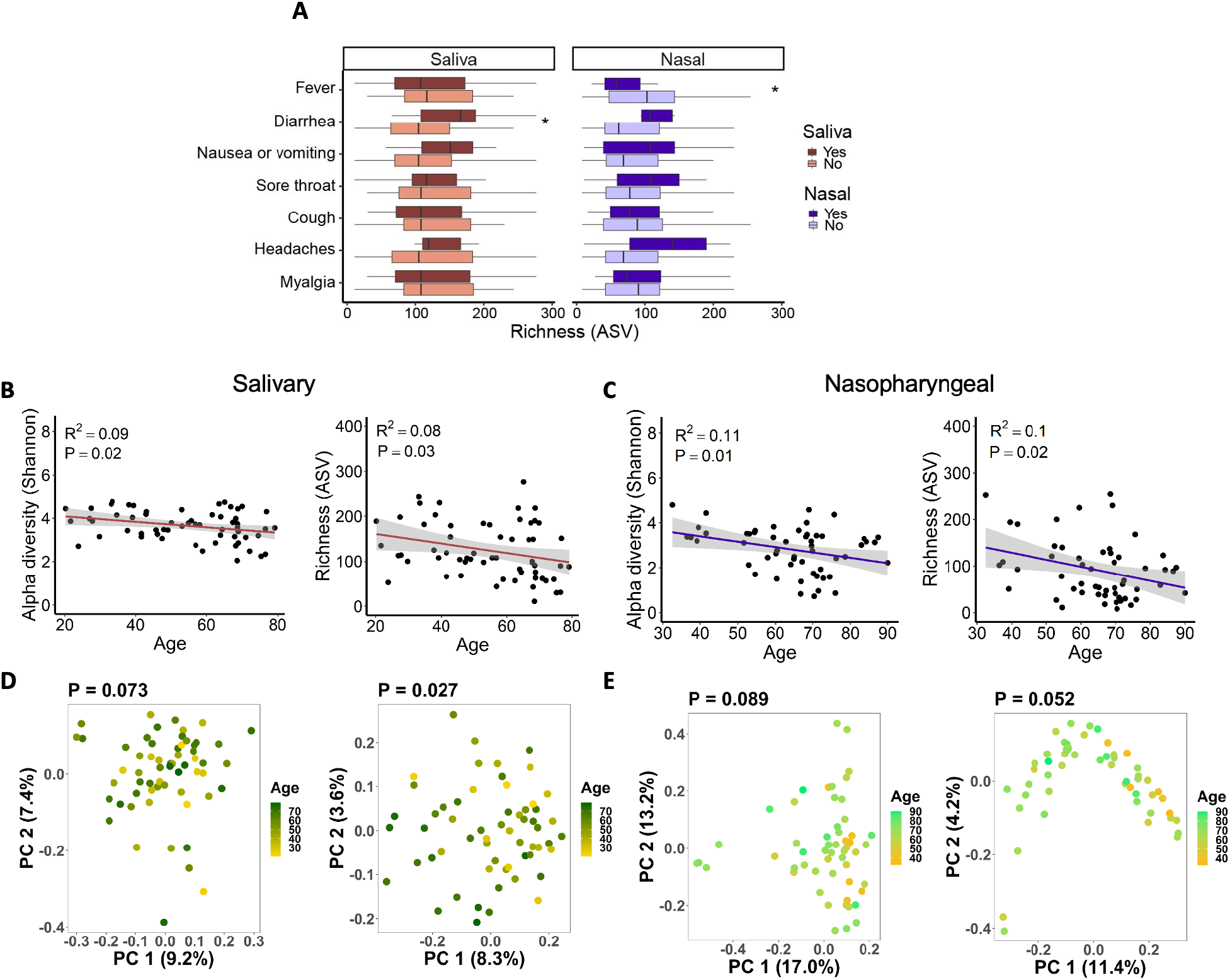
Associations between salivary and nasopharyngeal microbiomes of SARS-CoV-2-positive patients and COVID-19 symptoms and age. (A) Alpha diversity, represented by richness, in saliva (red) and nasopharyngeal swab (blue) samples, for patients with a given symptom (dark red or dark blue) or without (light red or light blue). Age versus alpha diversity represented by Shannon index and richness of salivary (B) and nasopharyngeal (C) microbial communities. Principal coordinates (PC) analysis of weighted (D, E, left) and unweighted (D, E, right) UniFrac distances for salivary (D) and nasopharyngeal (E) microbial communities with age. The shaded areas in panels B and C indicate the 95% confidence intervals. Statistical significance was assessed using the Wilcoxon signed-rank test for panel A, linear regression for panels B and C, and PERMANOVA for panels D and E. *, *P* < 0.05.

For both salivary and nasopharyngeal microbial communities, alpha diversity was significantly negatively correlated with age in patients with COVID-19 (**Figure 3B, Figure 3C**). Principal coordinates analysis of unweighted, but not weighted UniFrac distances, showed significant dissimilarity in the salivary microbiome by age (**Figure 3D**). A similar trend was observed in the nasopharyngeal microbiome but fell short of significance (*P* = 0.052, **Figure 3E**). Since unweighted UniFrac distances only evaluate differences in taxa between groups, unlike weighted UniFrac distances which also assess the abundance of each taxon, this result suggests that there are compositional differences with age, but the different bacterial taxa may have a relatively low abundance. These correlations did not achieve significance in patients without COVID-19, but the overall trends were similar (**Figure S3A–D**).

### 3. Several bacterial genera in the salivary and nasopharyngeal microbiome are differentially abundant between COVID-19 patients who were later admitted to an ICU and those who were not. Relative abundance of saliva *Bifidobacterium* is associated with plasma concentrations of IL-17F and MCP-1

To evaluate whether microbial differences by COVID-19 severity exist at the taxa level, we compared the salivary and nasopharyngeal microbiomes of ICU and non-ICU COVID-19 patients at the genus level using DESeq2 for differential abundance analysis. In saliva samples, three bacterial genera were significantly different between ICU and non-ICU groups, with *Bifidobacterium* (*P* = 0.00016), *Lactobacillus* (*P* = 0.0018), and *Solobacterium* (*P* = 0.026) being more abundant in the non-ICU group (**Figure 4A**). In nasopharyngeal samples, four genera were significantly different between the two groups, with *Paracoccus* (*P* = 0.0026) being more abundant in the non-ICU group and *Proteus* (*P* = 0.000036), *Cupravidus* (*P* = 0.023), *and Lactobacillus* (*P* = 0.023) being more abundant in the ICU group (**Figure 4B**). Differential abundance data is summarized in **Figure 4C**.

**Figure 4.**
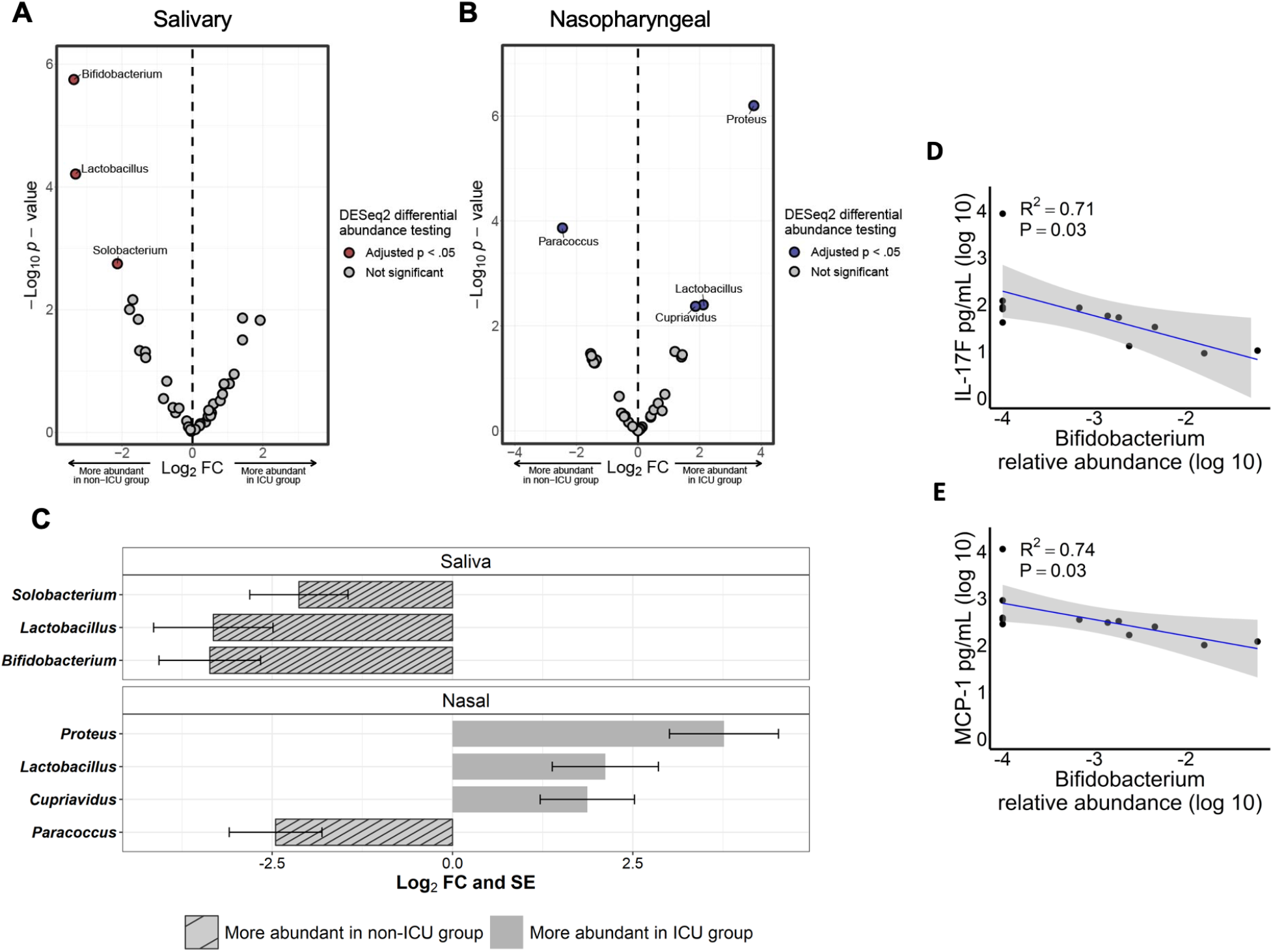
Differentially abundant bacterial genera in the salivary and nasopharyngeal microbiomes between ICU and non-ICU COVID-19 patients. Volcano plot of log_2_ fold change (FC) vs -log_10_ *P* value for salivary (A) and nasopharyngeal (B) microbial communities. Red and blue dots represent bacterial genera whose relative abundances were significantly different between the ICU and non-ICU groups. Significantly differentially abundant genera are summarized in panel C. The striped bars indicate genera that were more abundant in the non-ICU group while the solid bars indicate genera that were more abundant in the ICU group. Correlations were evaluated between salivary *Bifidobacterium* relative abundance (log 10) and plasma concentrations of IL-17F (D) and MCP-1 (E). DESeq2 *P* values were adjusted for multiple comparisons using the Benjamini-Hochberg method. The shaded areas in panels D and E indicate the 95% confidence intervals. Statistical significance was assessed in panels D and E using Spearman’s rank correlation coefficient.

Several studies have reported increased levels of inflammatory markers such as C-reactive protein (CRP), interleukin-2 (IL-2), IL-6, IL-7, IL-10, tumor necrosis factor alpha (TNF alpha), granulocyte colony-stimulating factor (G-CSF), and monocyte chemoattractant protein 1 (MCP-1) in COVID-19 patients with severe disease (24-26). To investigate whether the bacterial genera we identified as enriched or depleted in severe COVID-19 were correlated with systemic immune responses, we tested associations between the relative abundances of each genus and plasma concentrations of cytokines/blood markers in selected patients with COVID-19. We observed that a greater abundance of genus *Bifidobacterium* in the salivary microbiome was associated with lower levels of IL-17F and MCP-1 (**Figure 4D, Figure 4E**). Correlations between all profiled cytokines and bacterial genera enriched or depleted in severe COVID-19 cases are given in **Table S1** for saliva samples and **Table S2** for nasopharyngeal swab samples.

## Discussion

In this study, we profiled the salivary and nasopharyngeal microbiome of a COVID-19 cohort and validated that COVID-19 patients had significantly different microbial communities compared to those of non-COVID-19 patients. We then focused on the COVID-19 patients to identify microbial markers that are associated with disease severity. While there were no community level differences in the salivary and nasopharyngeal microbiomes of ICU and non-ICU groups, several bacterial genera including *Bifidobacterium, Lactobacillus, Solobacterium, Proteus, Cupriavidus*, and *Paracoccus*, correlated with COVID-19 severity. Salivary and nasopharyngeal microbiota also associated with COVID-19 symptoms, and relative abundance of *Bifidobacterium* in saliva was found to be associated with plasma concentrations of IL-17F and MCP-1.

Several studies have already characterized the oral microbiome in COVID-19 patients. In agreement with our results, multiple groups reported significant reductions in oral microbiome diversity in COVID-19 patients compared to non-COVID-19 controls (11, 20, 21). Multiple studies have also reported inverse correlations between oral microbiome alpha diversity and symptoms severity (11, 12, 21). One study identified a substantial decrease in alpha diversity in critical COVID-19 patients (defined as respiratory failure requiring mechanical ventilation, shock, or organ failure requiring ICU admission) compared to non-critical COVID-19 patients and healthy controls (8). The discrepancy between this result and our observations may suggest that classification of COVID-19 severity and sampling site are potential factors that may modify the correlations between microbiome and COVID-19. Few studies have performed differential abundance analysis of the oral microbiota between severe and non-severe COVID-19 patients. One study that utilized metagenomic sequencing of oropharyngeal swab samples from COVID-19 patients identified several species that were associated with COVID-19 severity, none of which were members of genera identified as associated with severity in our study (8). There could be several reasons for this discrepancy, including different patient demographics or differences in sequencing and analysis methods. More studies are needed to confirm these results and more confidently identify oral microbiota which may be helpful biomarkers of COVID-19 severity or modify host immune responses to impact viral progression.

So far, data has been mixed on the effect of COVID-19 on nasopharyngeal microbiota composition. Several early studies reported no major alterations in the nasopharyngeal microbiome after SARS-CoV-2 infection, while others reported substantial community level alterations to the nasopharyngeal microbiota after SARS-CoV-2 infection (22, 23, 27-29). Several studies have reported associations between nasopharyngeal microbiota and COVID-19 severity, with limited consistency in specific taxa associated with disease severity between studies (15, 30, 31). Recent evidence suggests that the contradictory results observed in COVID-19 respiratory microbiome studies may be driven by confounders such as time in ICU, oxygen support, and mechanical ventilation (32). These confounders may also explain discrepancies between our results and previous COVID-19 oral and airway microbiome studies.

We found *Bifidobacterium* was depleted in the salivary microbiome of ICU COVID-19 patients compared to non-ICU patients. This is consistent with a prior study which found depletion of *Bifidobacterium* in the oropharynx of ICU COVID-19 patients (15). Several studies have described the potential of *Bifidobacterium* to trigger immunomodulatory responses and maintain host physiological homeostasis (33-35). Mouse studies have also demonstrated the ability of oral and intranasally administered *Bifidobacterium* probiotics to protect against viral-induced lung inflammation and injury (36, 37). It has also been shown that certain strains of *Bifidobacterium* have the potential to suppress IL-17 production (38). Our study demonstrated a significant negative correlation between abundance of *Bifidobacterium* in the salivary microbiome and plasma levels of IL-17. The exact mechanisms by which IL-17 may contribute to inflammation and lung injury in SARS-CoV-2 infection are incompletely understood, but IL-17 response is known to mediate acute lung injury induced by viral infection (39-41). Furthermore, we found relative abundance of *Bifidobacterium* in the salivary microbiome was negatively correlated with plasma concentrations of MCP-1, another biomarker of severity in COVID-19 patients (42). Indeed, some strains of *Bifidobacterium* have been shown to downregulate MCP-1 levels in vitro and in vivo, suggesting an anti-inflammatory effect of certain *Bifidobacterium* strains (43, 44). Further work is needed to confirm these associations and elucidate any potential role of *Bifidobacterium* SARS-CoV-2 infection.

Interestingly, *Lactobacillus* was depleted in the salivary microbiome, but enriched in the nasopharyngeal microbiome of patients with severe COVID-19, indicating a potentially opposite, site-specific effect of *Lactobacillus* in disease outcome. Consistent with this hypothesis, previous studies showed that *Lactobacillus* in the gut microbiome was enriched in patients who recovered from COVID-19, while *Lactobacillus* in the upper respiratory tract microbiome was significantly associated with mortality in SARS-CoV-2-positive patients (19, 45). The immunomodulatory properties of *Lactobacillus* species are well described, and *Lactobacillus* strains have been widely used as probiotics. During viral respiratory tract infections, *Lactobacillus* can activate immune cells essential for antiviral defense and restrict viral replication (46, 47). In the context of SARS-CoV-2 infection, a recent study showed treatment of SARS-CoV-2 infected cells with *Lactobacillus plantarum* Probio-88 inhibited SARS-CoV-2 replication and production of reactive oxygen species and led to a reduction of inflammatory markers such as interferon alpha, interferon beta, and IL-6 (48). The antiviral activity of *Lactobacillus plantarum* may derive from their production of plantaricins, antiviral compounds with high binding affinity toward SARS-CoV-2 helicase that may prevent binding of ss-RNA during viral replication (48). Further study is needed to understand how *Lactobacillus* acts differently on the immune system in the upper airway compared to the gut and mouth.

We reported a correlation between the composition of the oral and upper respiratory microbiome with certain COVID-19 symptoms. The salivary microbiome of COVID-19 patients was associated with diarrhea, with COVID-19 patients with diarrhea having higher species abundance compared to those who did not have diarrhea. We also found that the nasal microbial community alpha diversity was significantly reduced in COVID-19 patients with fever than those without fever. While these results represent a potential link between oral and nasopharyngeal microbiota and COVID-19 pathophysiology, further research is needed to determine whether microbial dysbiosis predisposes the host to certain symptoms, if the observed microbial alterations are responses to patients’ symptoms and immune states, or if both respond to some other factor. In addition, we found that both salivary and nasopharyngeal microbiome alpha diversity negatively correlated with age in COVID-19 patients. The reduced diversity in salivary and nasopharyngeal bacterial species with aging could potentially predispose the elderly to severe COVID-19 (49).

Our study has several shortcomings which should be addressed. This study had a limited sample size and may be underpowered to detect certain differences between groups of interest. There were substantial differences in rates of hospitalization and co-morbidities such as congestive heart failure, diabetes, and hypertension, between the SARS-CoV-2-positive and SARS-CoV-2-negative groups, which could potentially impact our microbiome data. Although we included a matched case-control analysis, the sample size and power were greatly reduced. Our COVID-19 cohort included only symptomatic patients, and mildly symptomatic or asymptomatic individuals with COVID-19 may exhibit distinct microbiome features. The human microbiome can be influenced by diet, and information on diet was not collected in this study. Cytokine data was not available for all patients enrolled in the study, limiting the sample size for associations between the microbiome and systemic immune response. Furthermore, the human microbiome is highly variable across populations. In this study, all samples were collected in the greater St. Louis metropolitan area, potentially limiting its generalizability to the wider population.

In summary, we found several salivary and nasopharyngeal bacterial genera associated with COVID-19 severity. Although our findings cannot infer causality and should be validated in future studies with larger sample sies, this work provides additional information to characterize associations between COVID-19 and the human microbiome. This work may serve as a foundation for additional studies to uncover the underlying mechanisms linking the oral and airway microbiome to COVID-19 outcomes.

## Methods

### 1. Study participants

The patients included in this study were part of a prospective observational cohort of subjects with COVID-19-related symptoms who presented to Barnes-Jewish Hospital or affiliated Barnes-Jewish Hospital testing sites in Saint Louis, Missouri, USA, between March and September of 2020. Inclusion criteria required that subjects were symptomatic (fever, chills, conjunctival congestion, nasal congestion, headaches, cough, sputum production, sore throat, shortness of breath, nausea or vomiting, diarrhea, myalgia, fatigue, rash, lymphadenopathy, or confusion) and had a physician-ordered SARS-CoV-2 test performed in the course of their normal clinical care. Diagnosis of COVID-19 was based on a positive nasopharyngeal swab polymerase chain reaction test. Participants’ symptoms data were collected from surveys conducted when participants presented to a medical facility for testing and clinically relevant medical information such as ICU admission was collected from electronic medical records. This study was approved by the Institutional Review Board at Washington University in St. Louis (IRB number 202003085). All patients who were enrolled in the study provided informed consent prior to participation.

### 2. Sample collection, processing, and microbial DNA sequencing

Saliva and nasopharyngeal swab samples were collected at the time of enrollment, which was during or shortly following evaluation at a medical facility. The vast majority of samples were collected within 14 days of patients’ onset of COVID-19-related symptoms. For saliva collection, saliva was directly deposited into a container with an attached funnel and stored in a −80 degrees Celsius (°C) freezer until use. Nasopharyngeal swab samples were collected by a trained provider by inserting a swab along the nasal septum, just above the floor of the nasal passage, to the nasopharynx, until resistance was felt. Then, the swab was rotated several times before being withdrawn. Nasopharyngeal swabs were then placed in viral transport media and vortexed prior to being frozen at -80 degrees Celsius (°C) until use.

Prior to microbial DNA extraction, samples were heated at 56°C for 30 minutes to inactivate SARS-CoV-2 virus. Microbial DNA extraction of saliva and nasopharyngeal swab samples, sequencing library preparation, and 16S ribosomal RNA (rRNA) gene sequencing were performed as described previously (50). Briefly, genomic DNA was extracted from nasopharyngeal swab and saliva samples using the zymoBIOMICS DNA Miniprep Kit (Zymo Research, Irvine, California). 16S rRNA sequencing libraries were prepared by amplifying and barcoding the V1-V2 region of the 16S rRNA gene using the Quick-16S NGS Library Prep Kit (Zymo Research). Samples were pooled and sequenced on the Illumina MiSeq platform with 2 × 250 base pair standard run at Washington University DNA Sequencing Innovation Lab.

### 3. Sequencing data processing

Amplicon sequence variants (ASVs) were inferred from de-muliplexed fastq files using the DADA2 R package (https://benjjneb.github.io/dada2/tutorial.html) (51) and taxonomy was assigned from de-muliplexed fastq files using the Ribosomal Database Project’s Training Set 16. Sequencing data were quality filtered by trimming the last 10 nucleotides of each read to remove low quality tails then performing de-noising and chimera sequence removal using the default settings in the DADA2 pipeline. Statistical analyses were conducted in R version 3.4.2 and visualization was done with ggplot2 (https://ggplot2.tidyverse.org). Phyloseq, an R package (https://joey711.github.io/phyloseq/) (52), was used to calculate alpha diversity, beta diversity, and principal coordinates. To perform differential abundance testing, we used the R Package DESeq2 (https://bioconductor.org/packages/release/bioc/html/DESeq2.html) (53) which uses a generalized regression model with a logarithmic link, following a negative binomial distribution. DESeq2 *P* values were adjusted for multiple comparisons using the Benjamini-Hochberg method. Differential abundance analysis was conducted at all taxonomic levels and differentially abundant genera between the ICU and non-ICU groups identified by DESeq2 were displayed.

### 4. Cytokine quantification

Participant blood samples were collected within 24 hours of emergency department presentation in EDTA-containing vacutainers (BD Biosciences, San Jose, CA), transported on ice, spun down at 2500g for 10 min at 4°C, and stored at −80°C until further analysis. Cell-free plasma was analyzed using a human magnetic cytokine panel providing simultaneous measurement of 35 cytokines (Thermo Fisher Scientific, Waltham, MA). The assay was performed according to the manufacturer’s instructions with each subject sample performed in duplicate and then analyzed on a Luminex FLEXMAP 3D instrument.

### 5. Statistics

Differences between study groups were compared using the nonparametric Mann-Whitney U test for continuous variables and chi-squared test or Fisher’s exact test for categorical variables. Alpha diversity (Shannon index or observed species richness) differences between groups were compared using Wilcoxon signed-rank test. For beta diversity, principal coordinates analysis (PCoA) of weighted and unweighted UniFrac distances was performed to represent distances between microbial communities and differences in beta diversity between groups were evaluated using permutational multivariate analysis of variance (PERMANOVA) as implemented in the *adonis* function of the R package Vegan version 2.5-7. To evaluate correlations between blood marker concentrations and alpha diversity, we used linear regression. To evaluate correlations between blood marker concentrations and relative abundance of bacterial genera depleted or enriched in severe COVID-19 patients, we used Spearman’s rank correlation coefficient with false discovery rate (FDR) adjustment to correct for multiple comparisons. For all statistical tests, a *P* value of < 0.05 after controlling for multiple comparisons using the Benjamini-Hochberg method or FDR correction when appropriate, was considered to indicate significance.

## Author contributions

L.W. designed and coordinated this study. J.O., P.M., C.G. and A.R. established the cohort and collected samples from participants. J.K. processed samples for sequencing. A.Z. and L.W. performed sequencing data analysis. J.K. and L.W. wrote the manuscript. All authors provided critical comments on the manuscript and approved the manuscript.

## Acknowledgments

This work was supported by the National Institutes of Health [grant number 5R21AI139649-02, contact PI: Leyao Wang]. This study utilized samples obtained from the Washington University School of Medicine’s COVID-19 biorepository, which is supported by: the Barnes-Jewish Hospital Foundation; the Siteman Cancer Center grant P30 CA091842 from the National Cancer Institute of the National Institutes of Health; and the Washington University Institute of Clinical and Translational Sciences grant UL1TR002345 from the National Center for Advancing Translational Sciences of the National Institutes of Health. The content is solely the responsibility of the authors and does not necessarily represent the view of the NIH.

