## Supplementary material for "The salivary and nasopharyngeal microbiomes are associated with SARS-CoV-2 infection and disease severity"

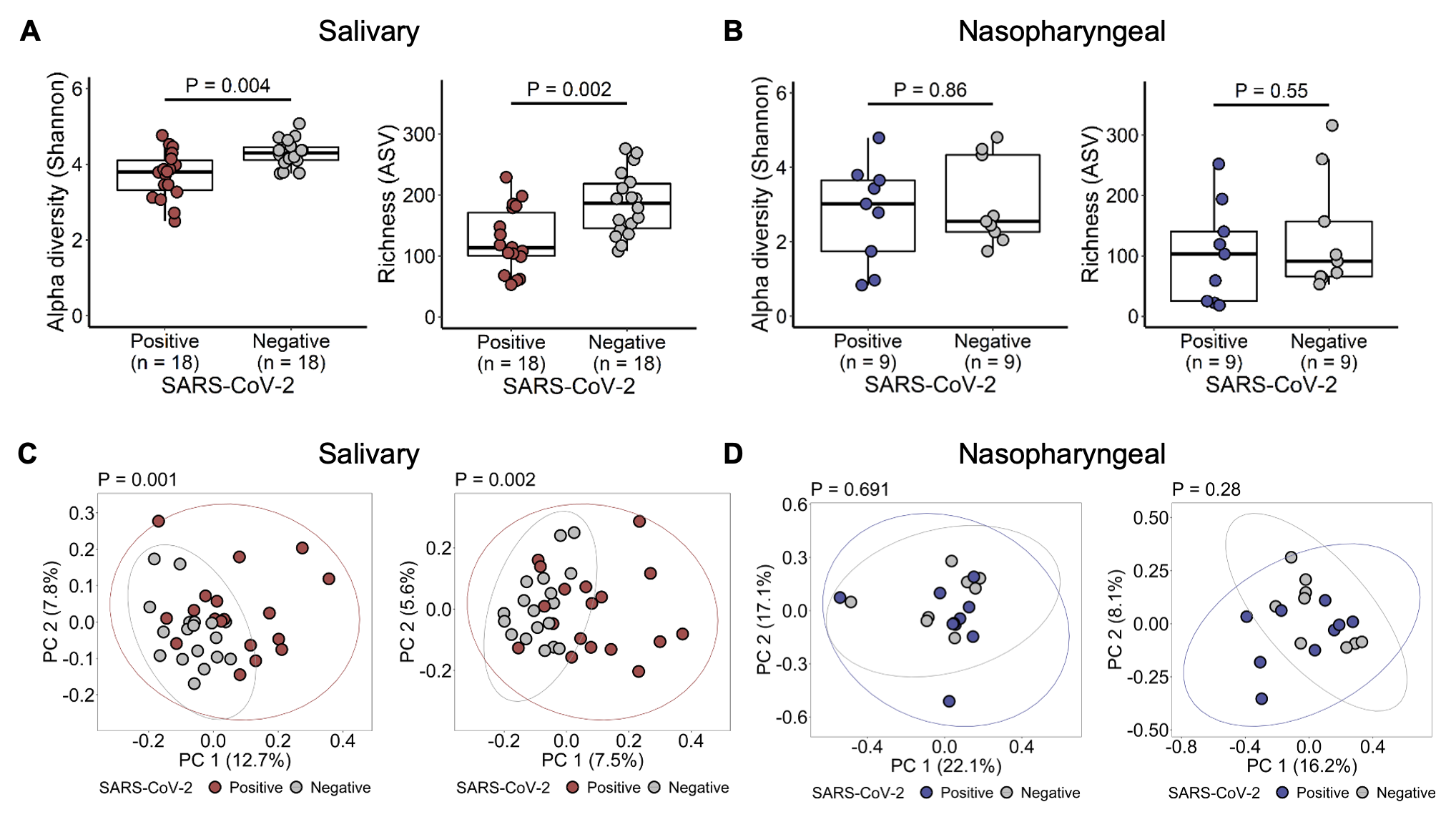

**Figure S1**. Community-level comparisons of the salivary and nasopharyngeal microbial communities between sex-, age-, and race-matched SARS-CoV-2-positive and SARS-CoV-2-negative patients. Alpha diversity was analyzed for salivary (A) and nasopharyngeal (B) microbial communities. Principal coordinates analysis of weighted and unweighted UniFrac distances was performed to compare the salivary (C) and nasopharyngeal (D) microbial communities of SARS-CoV-2-positive and SARS-CoV-2-negative subjects. Statistical significance was assessed using Wilcoxon signed-rank test for panels A and B, and using PERMANOVA for panels C and D.

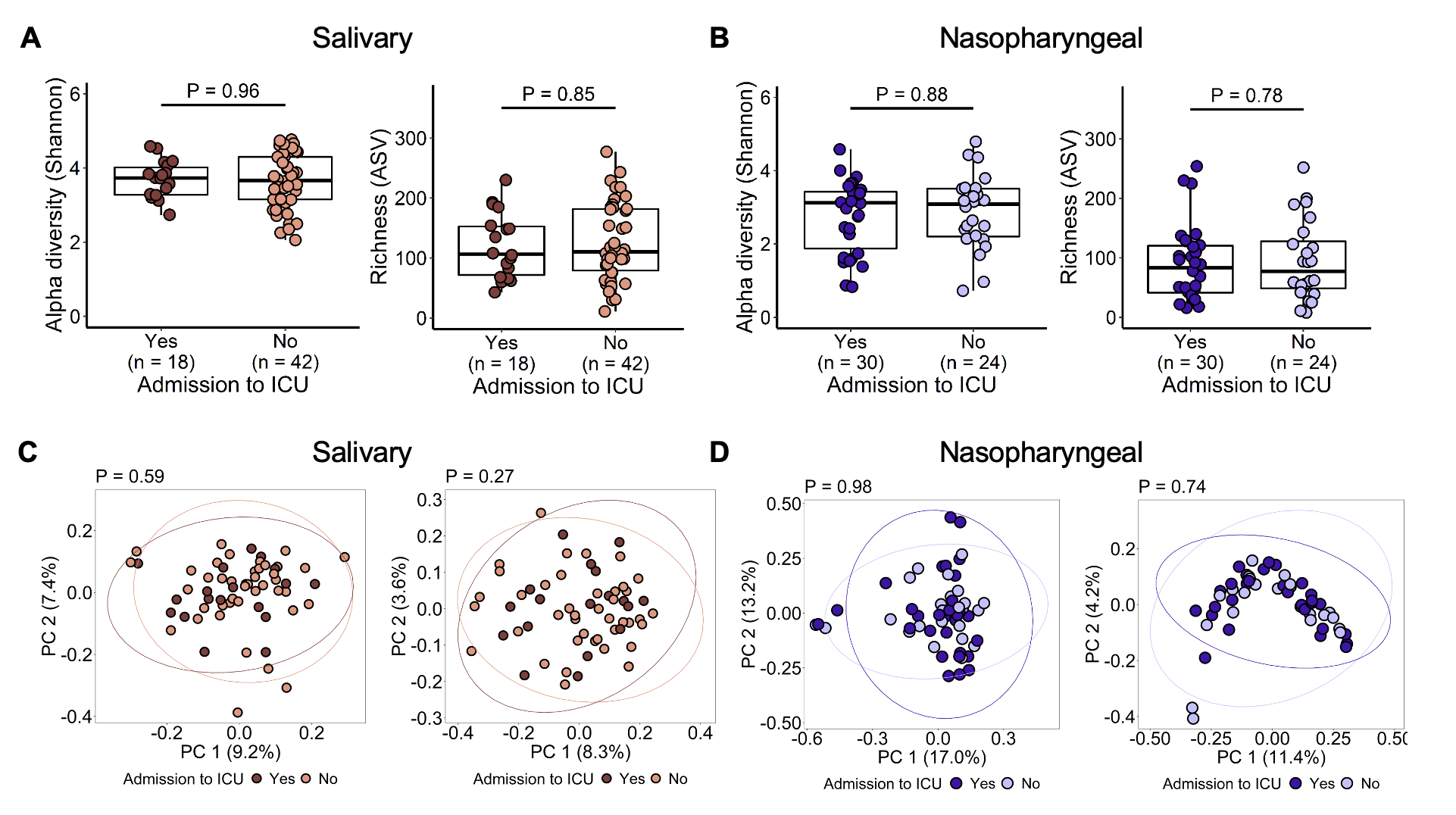

**Figure S2**. Community-level comparisons of the salivary and nasopharyngeal microbial communities between COVID-19 patients with or without severe disease. Alpha diversity was analyzed for salivary (A) and nasopharyngeal (B) microbial communities. Principal coordinates analysis of weighted and unweighted UniFrac distances was performed to compare the salivary (C) and nasopharyngeal (D) microbial communities of ICU and non-ICU subjects. Statistical significance was assessed using Wilcoxon signed-rank test for panels A and B, and using PERMANOVA for panels C and D.

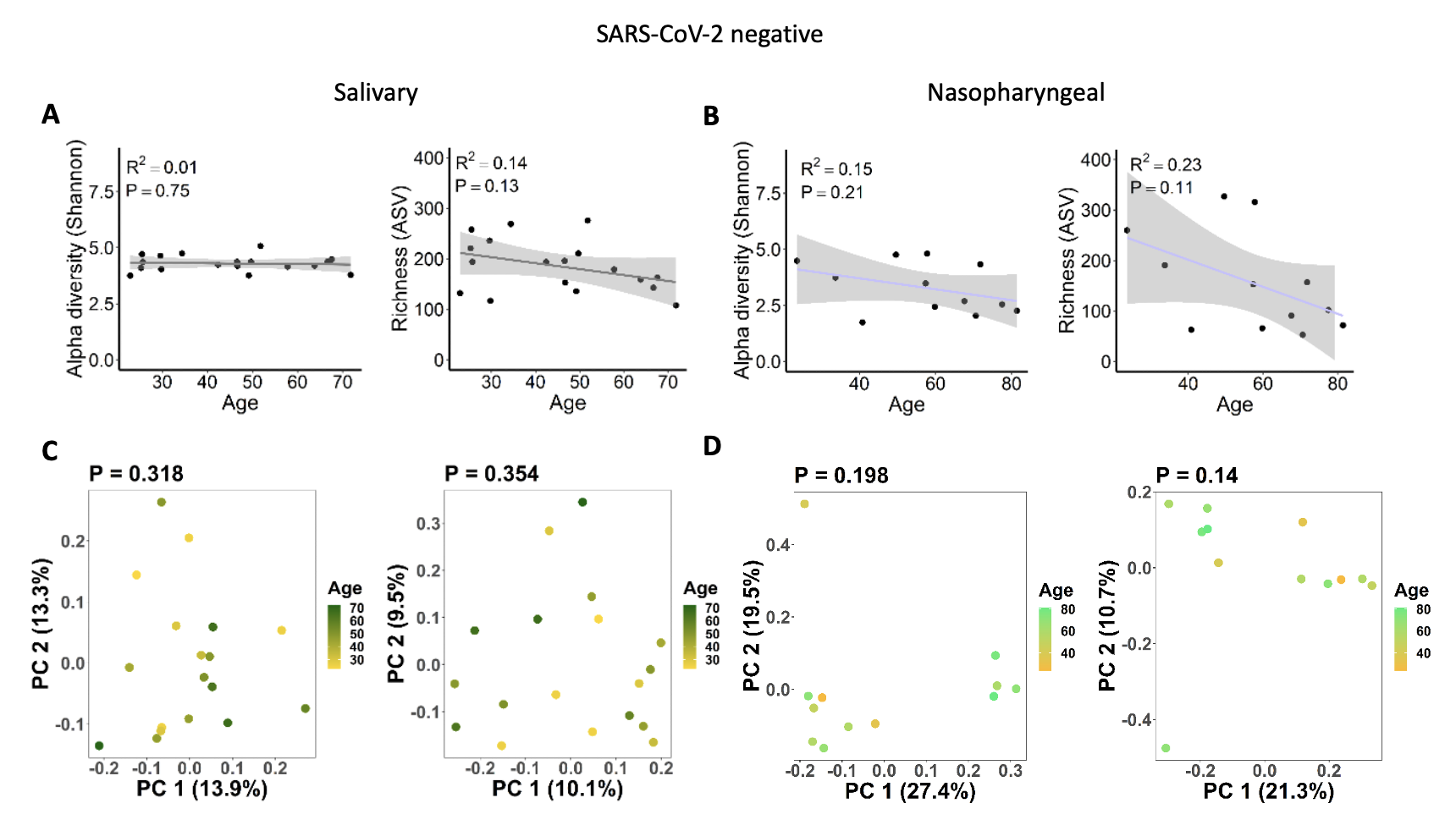

**Figure S3**. Associations between salivary and nasopharyngeal microbiomes of SARS-CoV-2 negative patients and age. Age versus alpha diversity represented by Shannon index and richness of salivary (A) and nasopharyngeal (B) microbial communities. Principal coordinates analysis of weighted (C, D, left) and unweighted (C, D, right) UniFrac distances for salivary (C) and nasopharyngeal (D) microbial communities. The shaded areas in panels A and B indicate the 95% confidence intervals. Statistical significance was assessed using linear regression for panels A and B, and PERMANOVA for panels C and D.
